# RexAB is essential for the mutagenic repair of *Staphylococcus aureus* DNA damage caused by co-trimoxazole

**DOI:** 10.1101/568196

**Authors:** Rebecca S. Clarke, Maya S. Bruderer, Kam Pou Ha, Andrew M. Edwards

**Affiliations:** MRC Centre for Molecular Bacteriology and Infection, Imperial College London, Armstrong Rd, London, SW7 2AZ, UK; Department of Infection Biology, Biozentrum, University of Basel, Klingelbergstrasse 70, CH-4056 Basel, Switzerland

**Keywords:** Reactive oxygen species, DNA damage, SOS response, *Staphylococcus aureus*

## Abstract

Co-trimoxazole (SXT) is a combination therapeutic that consists of sulfamethoxazole and trimethoprim that is increasingly used to treat skin and soft-tissue infections caused by methicillin-resistant *Staphylococcus aureus* (MRSA). However, the use of SXT is limited to the treatment of low-burden, superficial *S. aureus* infections and its therapeutic value is compromised by the frequent emergence of resistance. As a first step towards the identification of approaches to enhance the efficacy of SXT, we examined the role of bacterial DNA repair in antibiotic susceptibility and mutagenesis. This revealed that SXT caused DNA damage in *S. aureus* via both thymidine limitation and the generation of reactive oxygen species. Then, using mutants defective for DNA repair, it was found that repair of this damage required the RexAB nuclease/helicase complex, indicating that SXT causes DNA double-strand breaks. Furthermore, RexAB-mediated DNA repair led to induction of the SOS response, which resulted in an increased mutation rate and may explain the frequent emergence of resistant strains during SXT therapy. In summary, this work determined that SXT causes DNA damage in *S. aureus* via both thymidine limitation and oxidative stress, which is repaired by the RexAB complex, leading to induction of the mutagenic SOS response. Small molecule inhibitors of RexAB could therefore have therapeutic value by increasing the efficacy of SXT and decreasing the emergence of drug-resistance during treatment of infections caused by *S. aureus*.

## Introduction

*Staphylococcus aureus* is responsible for a wide spectrum of diseases ranging from superficial skin infections to life-threatening bacteraemia, endocartidis and toxic shock syndrome (1). Whilst β-lactam antibiotics such as oxacillin are first choice therapeutics for *S. aureus* infections, the prevalence of skin and soft tissue infections (SSTIs) caused by methicillin-resistant *S. aureus* (MRSA) strains has necessitated the use of second line therapeutics such as co-trimoxazole (SXT) (2, 3). SXT has several desirable properties including low cost, the availability of both oral and intravenous formulations and low host toxicity, making it an appealing treatment option (4).

SXT is a combination of two antibiotics; trimethoprim (TMP) and sulfamethoxazole (SMX) which target sequential steps in the tetrahydrofolate biosynthetic pathway (5). SMX inhibits dihydropteroate synthetase (DHPS), which prevents the production of dihydropteroic acid, while TMP binds and inhibits dihydrofolate reductase (DHFR), blocking the conversion of dihydrofolic acid to tetrahydrofolate (6). Since the production of tetrahydrofolate is essential for the biogenesis of thymidine, purines and some amino acids bacteria exposed to SXT will experience disrupted bacterial metabolism and stalled DNA replication (7).

Previous studies have reported that TMP-induced thymidine depletion contributes to bacterial cell death through stalled DNA replication forks, which together with the continued initiation of replication will result in DNA damage (8). However, one of the limitations of SXT as an antibiotic is that *S. aureus* can bypass SXT-mediated metabolic blockage by utilising exogenous thymidine released from damaged tissues (9). It is therefore hypothesised that the presence of thymidine at infection sites reduces the efficacy of SXT treatment in patients (10–12). As a consequence, SXT is only used for low burden superficial staphylococcal infections.

In addition to SXT-mediated DNA damage occurring via stalled replication forks, recent work with *E. coli* has implicated the production of reactive oxygen species (ROS) and maladaptive DNA repair in the bactericidal activity of this antibiotic (13). However, it is not clear if this is specific to *E. coli* or represents a general mechanism of bactericidal activity for many different bacteria.

SXT-mediated DNA damage triggers the SOS response, an inducible repair system that enables bacteria to survive genotoxic stress (14, 15). Regulation of the SOS response is highly conserved and occurs primarily via LexA, a transcriptional repressor of the SOS regulon (16). When DNA damage occurs, RecA binds to single-stranded DNA (ssDNA) at the lesion site to form a nucleoprotein filament which, i) facilitates DNA strand invasion that initiates repair by homologous recombination, and ii) induces the autocleavage of LexA, resulting in derepression of the SOS regulon (17). In *S. aureus*, it is believed that RecF assists RecA-binding to ssDNA-gaps from single-nucleotide lesions (18). However, double-strand breaks (DSB) are recognised by the AddAB complex (known as RexAB in *S. aureus*), which it binds and generates a ssDNA overhang through its helicase/exonuclease activity. This ssDNA strand then serves as the substrate for RecA-binding and activation of the SOS response (19).

The SOS-regulon of *S. aureus* consists of 16 LexA-regulated genes (17), including *recA*, *lexA* and *umuC*, which encodes a low-fidelity DNA polymerase. UmuC catalyses translesion DNA synthesis but lacks proofreading ability and permits DNA replication across unresolved lesions which often introduces mutations. Such mutagenic DNA repair may be advantageous to the pathogen by conferring resistance to antibiotics or adaptation to host stresses (17, 20). For example, mutations in the genes encoding DHFR and DHPS confer resistance to TMP and SMX respectively (21, 22). Therefore, exposure of *S. aureus* to SXT at concentrations that do not kill the pathogen may promote the emergence of resistance via induction of the mutagenic SOS response.

The prevalence of infections caused by drug-resistant pathogens necessitates new therapeutic options. However, given the lack of investment into the development of new antibiotics there is increasing interest in the development of therapeutics that enhance the efficacy of existing antibacterial drugs or by-pass resistance (23, 24). However, such an approach requires a thorough understanding of how antibiotics function and the mechanisms used by bacteria to survive exposure to antibacterial drugs. Therefore, the aims of this work were to identify the mechanisms by which SXT damages *S. aureus* DNA, the repair systems that this pathogen uses to withstand this damage and the consequences of repair for the emergence of resistance.

## Materials and Methods

### Bacterial strains and culture conditions

A full list of the bacterial strains used in this study is provided in Table 1. *S. aureus* was cultured in either Tryptic Soy Broth (TSB) or Mueller Hinton Broth (MHB) supplemented with calcium (25 mg/L) and magnesium (12.5 mg/L), to stationary phase for 18 h at 37 °C, with shaking (180 RPM). Culture media were supplemented with erythromycin (10 μg ml^-1^) for growth of transposon mutants, kanamycin (90 μg ml^-1^) for growth of the *recA* reporter strains, or anhydrotetracycline hydrochloride (AHT) (100 ng ml^-1^) to induce expression of *rexBA* on plasmids, as required (Table 1).

**Table 1.** Bacterial strains used in this study.

| Bacterial strain | Relevant characteristics | Source/ reference |
| --- | --- | --- |
| **Methicillin-resistant *S. aureus*** |  |  |
| USA300 LAC JE2 | LAC strain of the USA300 CA-MRSA lineage cured of plasmids | (53) |
| USA300 LAC JE2 *rexB*::Tn | USA300 LAC JE2 with a *bursa aurealis* transposon insertion in *rexB*, Ery<sup>r</sup> | (53) |
| USA300 LAC JE2 *umuC*::Tn | USA300 LAC JE2 with a *bursa aurealis* transposon insertion in *umuC*, Ery<sup>r</sup> | (53) |
| USA300 LAC JE2 *recF*::Tn | USA300 LAC JE2 with a *bursa aurealis* transposon insertion in *recF*, Ery<sup>r</sup> | (53) |
| USA300 LAC JE2 *dinB*::Tn | USA300 LAC JE2 with a *bursa aurealis* transposon insertion in *dinB*, Ery<sup>r</sup> | (53) |
| USA300 LAC JE2 *rexB*::Tn *pitet* empty | USA300 LAC JE2 with a *bursa aurealis* transposon insertion in *rexB* with integrated *pitet* empty plasmid, Ery<sup>r</sup> | This study |
| USA300 LAC JE2 *rexB*::Tn *pitet rexB* | USA300 LAC JE2 with a *bursa aurealis* transposon insertion in *rexB* with integrated *pitet* with AHT-inducible *rexB*, Ery<sup>r</sup> | This study |
| USA300 LAC JE2 +pCN34 *recA*-gfp | USA300 LAC JE2 containing pCN34 with *gfp* under the control of the *recA* promotor, Kan<sup>r</sup> | This study |
| USA300 LAC JE2 *rexB*::Tn +pCN34 *recA*-gfp | USA300 LAC JE2 with a *bursa aurealis* transposon insertion in *rexB* containing pCN34 with *gfp* under the control of the *recA* promotor, Kan<sup>r</sup> | This study |
| USA300 LAC JE2 +pCN34 Empty | USA300 LAC JE2 containing pCN34 empty vector, Kan<sup>r</sup> | This study |
| USA300 LAC JE2 *rexB*::Tn +pCN34 Empty | USA300 LAC JE2 with a *bursa aurealis* transposon insertion in *rexB* containing pCN34 empty vector, Kan<sup>r</sup> | This study |
| **Methicillin-sensitive *S. aureus*** |  |  |
| SH1000 | Methicillin sensitive *S. aureus* strain | (54) |
| SH1000 *rexB*::Tn | SH1000 with a *bursa aurealis* transposon insertion in *rexB*, Ery<sup>r</sup> | (20) |
| SH1000 *rexB*::Tn *pitet* empty | SH1000 with a *bursa aurealis* transposon insertion in *rexB* with integrated *pitet* empty plasmid, Ery<sup>r</sup> | This study |
| SH1000 *rexB*::Tn *pitet rexB* | SH1000 with a *bursa aurealis* transposon insertion in *rexB* with integrated *pitet* with AHT-inducible *rexB*, Ery<sup>r</sup> | This study |
| SH1000 +pCN34 *recA*-gfp | SH1000 containing pCN34 with *gfp* under the control of the *recA* promotor, Kan<sup>r</sup> | This study |
| SH1000 +pCN34 Empty | SH1000 containing pCN34 empty vector, Kan<sup>r</sup> | This study |

*S. aureus* SH1000 and JE2 mutants deficient for *rexB* were complemented using p*itet* (25) containing the wildtype *rexBA* operon. Since insertion of the transposon into *rexB* is expected to block the expression of the downstream *rexA* gene, the *rexB* mutants were complemented with the entire *rexBA* operon. The p*itet* vector is a single-copy plasmid that integrates into the *geh* locus of the *S. aureus* genome. It contains a tetracycline-inducible promoter that was used to control *rexBA* gene expression. Primers containing the AvrII or PmeI restriction sites (Table 2) were used to amplify the *rexBA* operon from USA300 JE2 genomic DNA. The PCR product and p*itet* vector were digested using AvrII and PmeI restriction enzymes, ligated using T4 ligase and transformed into CaCl_2_-competent *E. coli* DH5α cells. Successful ligation of *rexBA* into p*itet* was confirmed with DNA sequencing. Plasmid DNA was subsequently transformed into the DC10B *E. coli* strain which lacks cytosine methylation to allow bypassing of the staphylococcal restriction-modification barrier (26) and allowed successful electroporation into electrocompetent S*. aureus rexB* mutants in the JE2 and SH1000 genetic backgrounds. Empty vector that did not contain the *rexBA* operon was used as a control. Successful integration into the *S. aureus* genome was confirmed by PCR amplification of the *geh* gene followed by DNA sequencing.

**Table 2.**
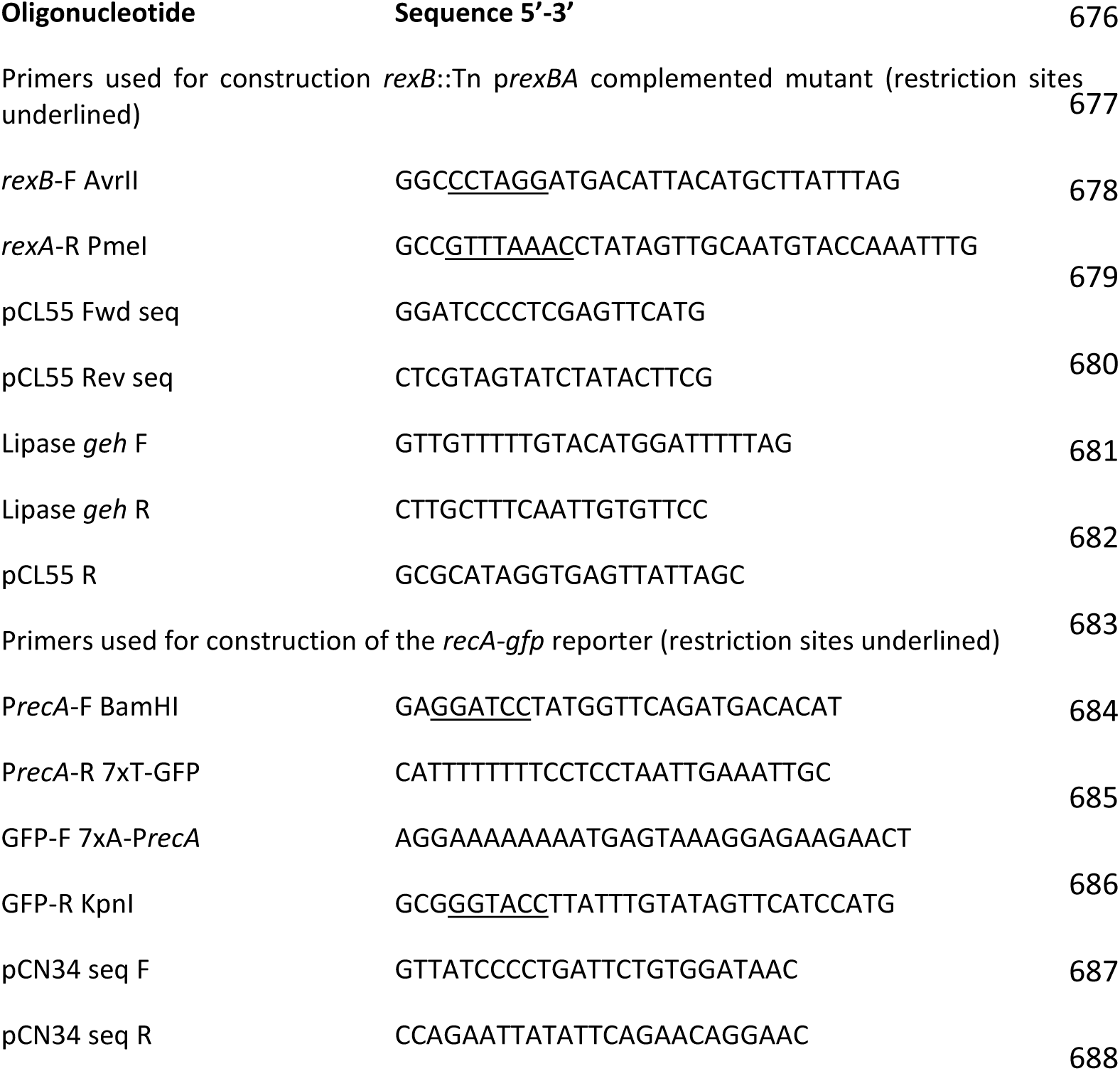
Primers used for construction of complemented *rexB* mutants and *recA*-gfp reporter.

### Minimum Inhibitory Concentration (MIC) Assay

The minimum inhibitory concentration was determined according to the broth microdilution protocol described previously (27). Briefly, 95 µl aliquots of either MHB or TSB supplemented with doubling concentrations of Co-trimoxazole (SXT), Trimethoprim (TMP) or Sulfamethoxazole (SMX) ranging from 0.008 μg ml^-1^ to 512 μg ml^-1^ were prepared in wells of 96 well plates. In this study, the concentration of SXT was calculated based on the concentration of TMP present within the combination therapeutic. Stationary phase cultures of *S. aureus* were diluted and inoculated into the antibiotic-containing media to a final cell density of 1 x 10^5^ CFU ml^-1^. For complemented mutants containing *rexBA* under the control of the p*itet* promoter, the medium was supplemented with AHT (100 ng ml^-1^) to induce gene expression. Plates were incubated for 17 h at 37 °C before OD_600_ readings were taken. The MIC was determined as the lowest concentration of antibiotic required to cause growth inhibition. To determine MICs under anaerobic conditions, 96 well plates were incubated in anaerobic jars containing Anaerogen gas packs at 37 °C for 18 h.

### SXT Killing Assay

Stationary phase *S. aureus* cultures were washed twice by alternate rounds of centrifugation and resuspension in fresh MHB or TSB (3 ml) to a density of 2 x 10^8^ CFU ml^-1^ containing 4 μg ml^-1^ SXT, the antibiotic clinical breakpoint concentration (28). Bacteria were incubated with SXT at 37°C with shaking (180 RPM) and survival determined over time by quantification of CFUs using the Miles and Misra method, which consists of serial dilution followed by plating onto agar plates to enable the enumeration of colonies (29). Data were converted to % survival by comparing CFU counts at specific time points with the starting inoculum. To investigate SXT-mediated killing under anaerobic conditions, washed stationary phase cultures were inoculated into pre-reduced broth in bijoux tubes supplemented with 4 μg ml^-1^ SXT, and incubated statically at 37 °C with the vessel caps loosened in an anaerobic jar containing an Anaerogen gas pack. The growth of *S. aureus* strains in TSB, under aerobic or anaerobic conditions was examined using a similar protocol using TSB or MHB without antibiotics.

### Disc-diffusion Assay

Stationary phase *S. aureus* cultures were adjusted to an OD_600_ of 0.063 (0.5 McFarland) in TSB and used to inoculate TSA plates using a cotton wool swab. Antibiotic discs containing 2.5 μg of either SXT or TMP (Oxoid) were placed onto the inoculated agar and incubated at 37 °C under aerobic or anaerobic conditions and incubated at 37 °C for 17 h. The zone of inhibition was measured as the total diameter of the cleared bacterial lawn, minus the diameter of the antibiotic disc.

### Mutation rate analyses

Mutation rates were determined as described previously (20, 30). Briefly, 30 parallel cultures of *S. aureus* were used for each condition or strain in bijoux tubes containing 1 ml TSB inoculated with *S. aureus* at ~ 5 x 10^5^ CFU ml^-1^ and incubated with or without 0.05 μg ml^-1^ or 0.1 μg ml^-1^ SXT with shaking at 37 °C for 24 h. After incubation, 10 cultures were selected at random, and total CFU counts determined through serial dilution in PBS followed by plating onto TSA plates. Subsequently, 100 μl of each of the 30 undiluted cultures were spread on TSA plates containing rifampicin (100 μg ml^-1^) before incubation at 37 °C for 24 h. The number of rifampicin-resistant colonies on each plate were counted, and the mutation rate with 95% confidence intervals calculated using the maximum-likelihood setting of the FALCOR mutation rate calculator (30). Since these assays were done in the presence of SXT it was important to use rifampicin resistance (rather than SXT resistance) to measure the mutation rate to avoid selecting for resistance emergence. Rifampicin resistance is a well-established marker for mutation-rate analysis and is not selected for or against under the conditions used.

### *recA*-reporter assay

Induction of the SOS response in *S. aureus* in response to SXT exposure was determined through the use of strains containing a *recA-gfp* reporter construct. These were generated by transforming *S. aureus* with pCN34 containing the *recA* promoter upstream of *gfp*. Primers, detailed in Table 2, were used to amplify the *recA* promoter region (sequence detailed in (14) from JE2 wildtype genomic DNA, and the *gfp* gene from the pCL55 P3 GFP plasmid (31). These products were combined by overlapping extension PCR using primers ‘PrecA-F BamHI’ and ‘GFP-R KpnI’, digested with BamHI and KpnI (relevant restriction sites incorporated into amplicon primers), and ligated into phosphatase-treated, BamHI and KpnI digested low copy shuttle vector pCN34. Empty vector without the *recA-gfp* construct served as a control. The *recA-gfp* pCN34 vectors were constructed in *E. coli* DC10B and confirmed by sequencing. Cultures of DC10B containing the *recA-gfp* pCN34 vector and empty vector were cultured in LB broth supplemented with ampicillin (100 μg/ml) to select for plasmid maintenance. Using methods described in (26), *recA-gfp* and empty vectors were transformed directly with into *S. aureus* WT JE2 and the *rexB*::Tn mutant. Reporter strains were grown in the presence of 90 μg/ml of kanamycin to maintain the plasmid.

Stationary phase cultures of *recA-gfp* reporter strains were pelleted by centrifugation, washed twice, and resuspended in TSB supplemented with kanamycin (90 μg ml^-1^). Cultures at ~ 3.33 x 10^8^ CFU ml^-1^ were exposed to a range of SXT concentrations (0.01 μg m^l-1^ to 8 μg ml^-1^) in black-walled microtiter plates. These were incubated, with shaking, at 37 °C for 17 h in a TECAN Infinite® 200 PRO microplate reader, where OD_600_ and GFP relative fluorescent units (RFU) (Excitation: 475 nm; Emission: 525 nm) was measured every 1000 seconds. The values were blanked against values generated from uninoculated wells, and GFP fluorescence was normalised to OD_600_ readings to determine *recA* expression levels.

### Endogenous ROS production

Production of ROS was detected using the cell permanent 2’,7’-dichlorodihydrofluorescein diacetate dye (H_2_DCFDA), which is converted to fluorescent DCF by oxidative cleavage of acetate groups (32). A 96 well microtitre plate assay was prepared in a similar format to the *recA-gfp* reporter assay, with the addition of 25 μM H_2_DCFDA and various concentrations of SXT. Wildtype and *rexB*::Tn mutant strains were washed and inoculated at ~ 3.33 x 10^8^ CFU ml^-1^. Plates were incubated with shaking at 37 °C as described previously, and OD_600_ and fluorescence (Excitation: 495 nm; Emission: 525nm) was measured every 1000 s. OD_600_ values were blanked against uninoculated wells, and fluorescence data was blanked to untreated wells. Fluorescent values were normalised to OD_600_ readings to determine production of ROS relative to cell number.

### Statistical Analysis

Means or medians were calculated from at least three biologically independent replicates and analysed by Student’s t test (two tailed, unpaired, assuming equal variances), One-way ANOVA or Two-way ANOVA corrected for multiple comparison using GraphPad Prism (V7.0) as described in the figure legends. MIC-bar graphs show the median value. All remaining graphs were plotted to show mean ± S.E.M. Error bars were omitted on *recA*-reporter data for clarity.

## Results

### Inactivation of *rexB* sensitises *S. aureus* to SXT

To investigate the role of bacterial DNA repair in the susceptibility of *S. aureus* to SXT, a panel of *S. aureus* transposon mutants were subjected to MIC and disc diffusion susceptibility tests, including SXT and its constituent antibiotics, TMP and SMX. These mutants lacked components of DNA repair (*rexA, rexB*, *recF*, *umuC* and *dinB*) previously shown to contribute to the repair of damage caused by the quinolone antibiotic ciprofloxacin, or acting as controls (20). Mutants in two distinct genetic backgrounds were used in this study; SH1000, a well-established laboratory strain, and JE2, a community-associated methicillin-resistant *S. aureus* (CA-MRSA) strain of the USA300 lineage, responsible for severe skin infections that are often treated with SXT (3, 33).

The MIC values for both the SH1000 and JE2 wild-type (WT) strains indicated susceptibility to SXT and TMP (< 2 µg ml^-1^), but resistance to SMX (>512 µg ml^-1^) based on EUCAST breakpoints for *S. aureus* [EUCAST, 2019] (Fig. 1A). Disruption of *rexB* in both JE2 and SH1000 strains lowered the MIC of SXT and TMP by 2-4-fold and 16-fold for SMX, whilst complementation of the *rexB* mutant either fully or partially restored wild-type levels of susceptibility to each of the antibiotics (Fig. 1A). The mutant lacking *recF* in the JE2 strain was also more susceptible than the WT to all three antibiotics, but the disruption of *dinB* or *umuC* only resulted in a slightly increased susceptibility to SXT but not TMP or SMX.

**Figure 1.**
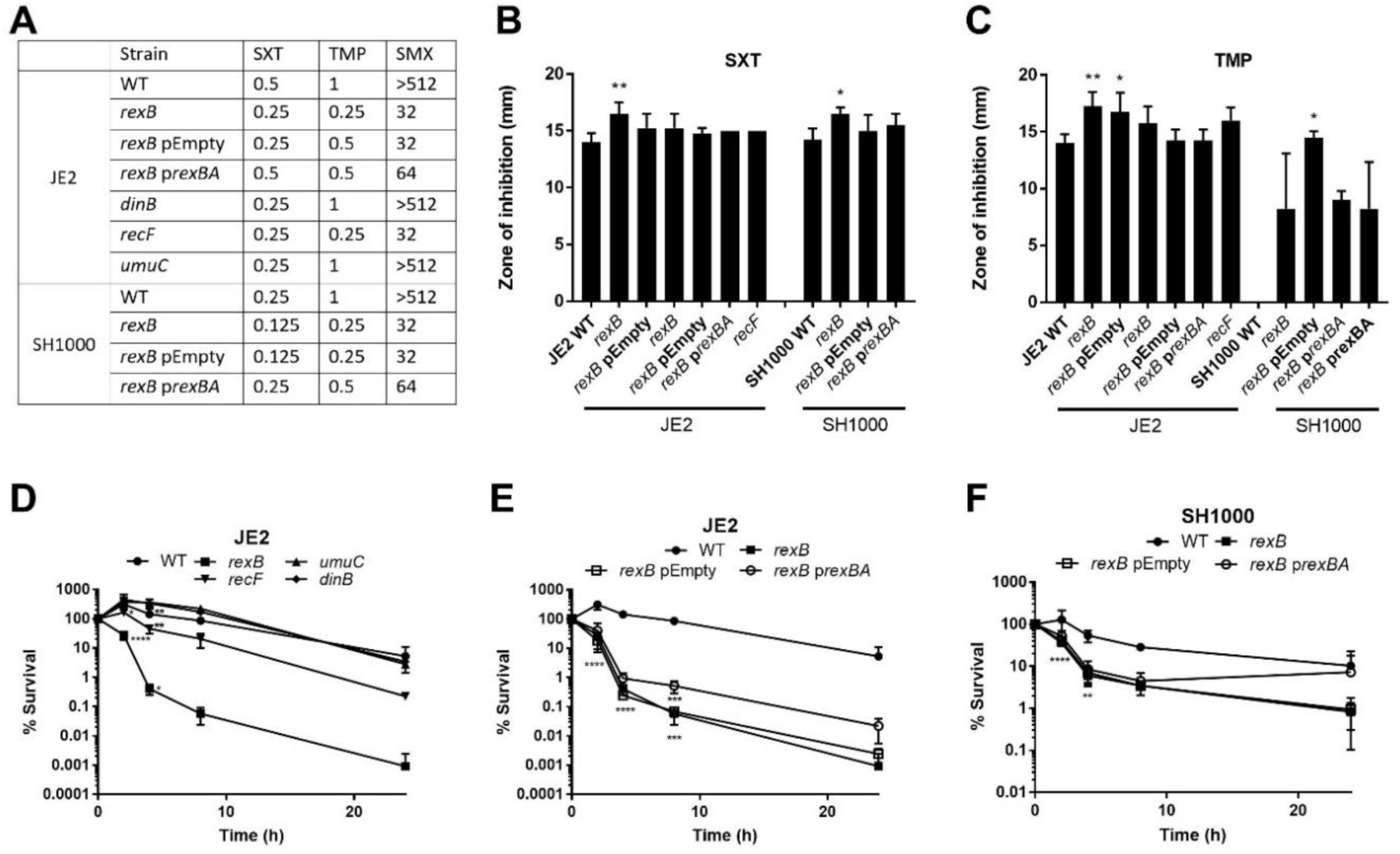
Mutants defective for DNA repair have increased susceptibility to SXT and its constituent antibiotics TMP and SMX. (A) MIC values for SXT, TMP and SMX for wild-type *S. aureus* JE2 and SH1000 and mutants defective for DNA damage repair in TSB. Data represent median values from n=3. (B,C) Zones of growth inhibition of WT JE2 and SH1000 *S. aureus* strains or damage-repair mutants from paper discs impregnated with 2.5 μg SXT (B) or TMP (C) after 16 h of incubation on TSA plates. Graphs represents mean±s.e.m from n=4. Values that are significantly different from the WT within each antibiotic-exposure were identified by one-way ANOVA). (D,E,F) Time-course survival assays of *S. aureus* JE2 (D,E) and SH1000 (F) and their derived DNA damage-repair mutants during incubation in TSB supplemented with 4 μg ml^-1^ SXT. Data are split between D and E for clarity. Percentage survival at each time point was calculated relative to the starting inoculum. Graphs represent mean±s.e.m from n=3. Values that are significantly different from the WT were identified by two-way ANOVA corrected for multiple comparison using the Dunnett method. P *≤* 0.05(*), P *≤* 0.01(**), P *≤* 0.001(***), P *≤* 0.0001(****).

Consistent with the MIC data, disc diffusion assays with both JE2 and SH100 strains showed that inactivation of *rexB* significantly increased the susceptibility of *S. aureus* to SXT and TMP, relative to the WT (Fig. 1B,C). Complementation of the *rexB* mutant with (p*rexBA*) reduced susceptibility close to WT levels, confirming the role of the RexAB complex (Fig. 1B,C). By contrast, inactivation of *dinB or umuC* did not significantly increase zone sizes for either antibiotic (Fig. 1B,C). However, the *recF*-deficient mutant showed an increase in susceptibility to TMP in the JE2 strain (Fig. 1C), indicating that both *rexBA* and *recF* contribute to the repair of DNA damage caused by TMP.

The finding that RexAB and RecF reduced the susceptibility of *S. aureus* to SXT confirmed that DNA repair modulated susceptibility to SXT and prompted us to undertake a screen of mutants with disrupted copies of *recJ*, *uvrA*, *uvrB*, *nth*, *mutS*, *mutL*, *sbcC* or *sbcD*, to identify any additional repair processes that might be involved. However, none of these mutants had a SXT MIC that was different from that of the wild-type.

Having established that staphylococcal DNA repair reduced the growth-inhibitory activity of SXT, we next investigated its role in the survival of *S. aureus* exposed to the breakpoint concentration of the antibiotic (4 μg ml^-1^) (EUCAST, 2019). There was a 20-fold reduction in survival over 24 h for the JE2 strain (Fig. 1D), indicating a relatively high level of tolerance to SXT at this concentration. Similar levels of survival were seen for mutants lacking *umuC* or *dinB* in the JE2 background (Fig. 1D). By contrast, survival of the *recF* mutant was reduced by >20-fold, relative to the WT, indicating increased susceptibility. Most notably however, the *rexB* mutant showed >5000-fold reduction in CFU counts relative to the WT at 24 h. Complementation of the *rexB* mutant increased survival 10-fold relative to a plasmid only control (Fig. 1E). Subsequent experiments with the SH1000 confirmed a role for *rexBA* in modulating susceptibility of *S. aureus* to SXT. There was a 12-fold greater loss of CFU counts of the *rexB* mutant relative to WT (Fig. 1F). However, similar to what was seen for the MIC assays, complementation of the *rexB* mutant with p*rexBA* restored survival levels close to those of the WT (Fig. 1F). Combined, these findings indicate that SXT causes DNA double-strand breaks in *S. aureus*, the repair of which by RexAB significantly promotes staphylococcal survival during exposure to the antibiotic.

### Thymidine limitation contributes to SXT-mediated DNA damage

Inhibition of folate biosynthesis by SXT results in abrogation of endogenous thymidine biosynthesis (34), which could provide an explanation for the DNA damage phenotype described above (Fig. 1). In support of this, previous studies have reported that the *in vitro* inhibitory activity of TMP and SXT is inversely proportional to the availability of thymidine in the culture medium (10, 35). The presence of exogenous thymidine allows *S. aureus* to continue with canonical DNA synthesis and allow effective DNA repair to occur, despite SXT-mediated inhibition of tetrahydrofolate production (13).

The experiments detailed thus far were performed in Tryptic Soy Broth (TSB), which contains a low concentration of thymidine (0.25 µM, (35)). To investigate the influence of exogenous thymidine on SXT activity in our assays, the growth of WT and *rexB* mutant strains was determined in TSB supplemented with doubling dilutions of SXT and thymidine via chequerboard assays (Fig. 2A-D). As expected, the presence of high concentrations of thymidine enabled growth of WT strains of both JE2 and SH1000 at higher concentrations of SXT (35) (Fig. 2A,C).

**Figure 2.**
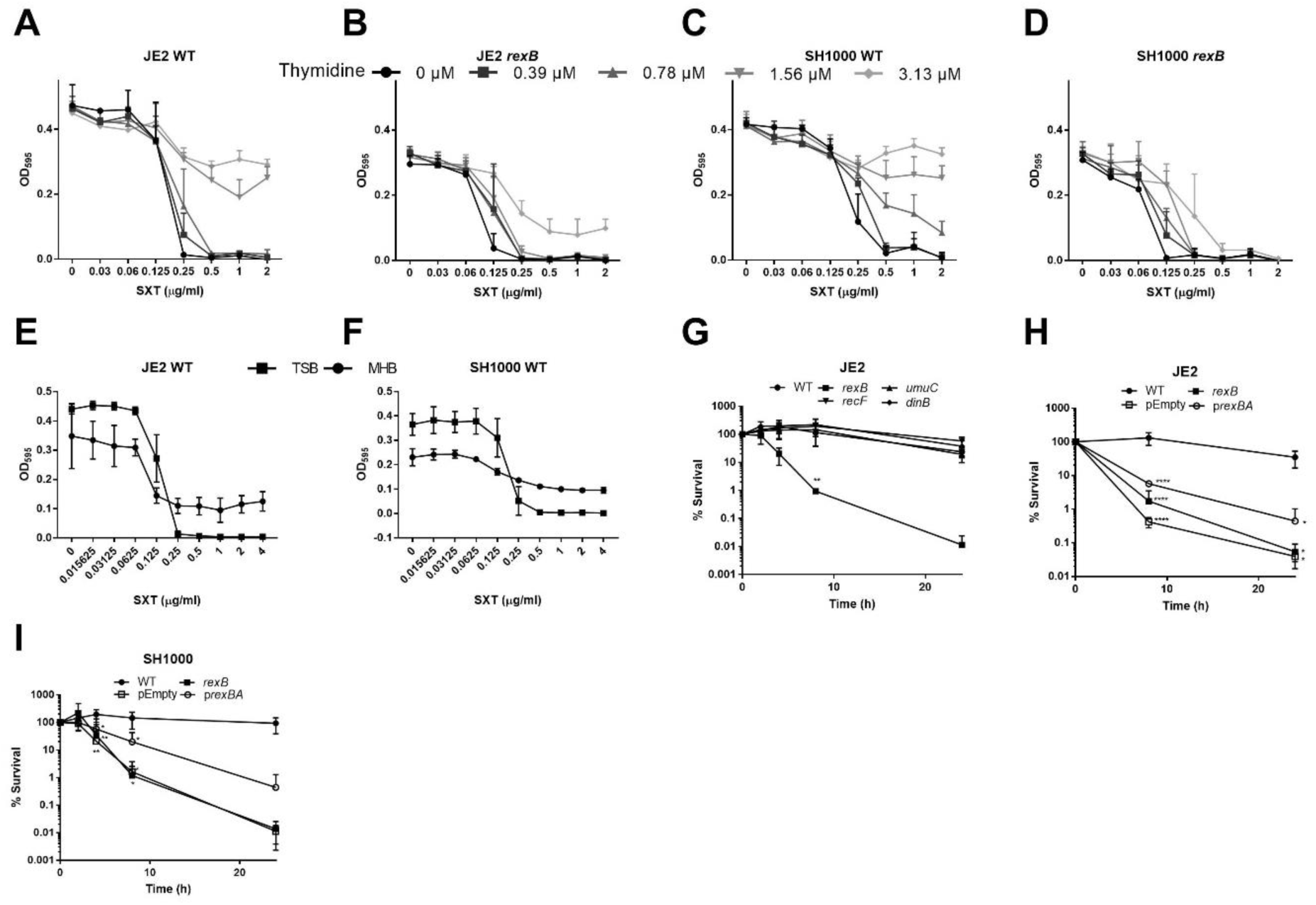
Mutants defective for *rexB* have increased susceptibility to SXT under thymidine-rich conditions. (A-D) Growth (OD_600_) of *S. aureus* JE2 WT (A) a JE2-derived *rexB* mutant (B), WT SH1000 (C) and SH1000-derived *rexB* mutant (D) in the presence of doubling concentrations of SXT in TSB supplemented with 0 - 3.13 μM thymidine as indicated. Graphs represent mean±s.e.m from n=3. (E-F) Growth (OD_595_) of S*. aureus* JE2 WT (E) and WT SH1000 (F) in TSB and MHB supplemented with doubling dilution concentrations of SXT. Graphs represent mean±s.e.m from n=3. (G-I) Time-course survival assays of *S. aureus* JE2 WT or mutants defective for DNA damage repair (G), *S. aureus* JE2 WT, *rexB* mutant only, or transformed with empty vector (pEmpty) or complemented mutant (p*rexBA*) (H) and SH1000 WT, *rexB* mutant only and *rexB* mutant transformed with pEmpty or p*rexBA* (I). For (G-I) all strains were incubated in MHB containing 4 μg ml^-1^ SXT. Graphs represent mean±s.e.m from n=3. Values that are significantly different from the WT were determined by two-way ANOVA corrected for multiple comparison using the Dunnett method. P *≤* 0.05(*), P *≤* 0.01(**), P *≤* 0.001(***), P *≤* 0.0001(****).

Given the thymidine-dependent activity of SXT, different culture media compositions may greatly affect antibiotic susceptibility (10, 35). For example, Mueller-Hinton Broth (MHB), the gold standard medium for antibiotic susceptibility testing, contains >16-fold more thymidine than TSB (35). Therefore, similar susceptibility tests were undertaken using this medium. Consistent with the checkerboard data, the efficacy of SXT in MHB was reduced relative to TSB, with detectable bacterial growth of both WT strains observed even at high concentrations of SXT, most likely due to the increased levels of thymidine (Fig. 2E,F).

To explore the effect of exogenous thymidine on the bactericidal activity of SXT, time-course killing assays were undertaken using MHB. As expected from the MIC assays, and by contrast to assays done in TSB (Fig. 1D) there was almost no killing of the *S. aureus* JE2 WT strain by SXT (4 μg ml^-1^) in MHB over 24 h (Fig. 2G). Similarly high levels of survival were observed for the *dinB*, *umuC* and *recF* mutants (Fig. 2G). By contrast, survival of the *rexB* mutant in MHB containing SXT was >2000-fold lower than that of the WT, despite the presence of exogenous thymidine (Fig. 2G). As for TSB, complementation of the *rexB* mutant (p*rexBA*) significantly promoted survival in MHB containing SXT, relative to a plasmid only control (Fig. 2H). Similar results were seen for the SH1000 strain, with the WT surviving in MHB containing SXT (4 μg ml^-1^), whilst viability of the *rexB* mutant was reduced >6000-fold relative to the inoculum (Fig. 2I). As for the JE2 strain, complementation of the SH1000 *rexB* mutant significantly promoted survival (> 10-fold) (Fig. 2I).

Combined, these data demonstrate that the RexAB DNA repair complex is required for the survival of *S. aureus* exposed to SXT, even when thymidine is available, suggesting that thymidine limitation is not the only mechanism by which this antibiotic damages bacterial DNA.

### Inactivation of *rexB* sensitises *S. aureus* to SXT-mediated growth inhibition but not killing under anaerobic conditions

It was recently reported that TMP exposure increases the production of reactive oxygen species (ROS) by *E. coli*, which together with incomplete DNA repair contributes to bacterial cell death (13). Therefore, we hypothesised that SXT-mediated DNA damage could occur via stalled DNA synthesis caused by inhibition of thymidine production and via the generation of ROS.

To examine whether the production of ROS contributes to SXT activity against *S. aureus*, the susceptibility of WT and DNA repair mutants to the antibiotic was determined under both aerobic and anaerobic conditions in low-thymidine TSB. In the absence of oxygen, the MIC of WT *S. aureus* was increased 4-fold for both JE2 and SH1000 strains (Fig. 3A). By contrast, the SXT MIC of the JE2 *rexB* mutant only doubled under anaerobic conditions, whilst the MIC of the SH1000 *rexB* mutant was unaffected by the presence of oxygen (Fig. 3A). To explore this further, zone of inhibition assays were done under aerobic and anaerobic conditions. Zones of inhibition were significantly reduced in the absence of oxygen for all strains and mutants, implying reduced susceptibility (Fig. 3B). However, despite the reduced antibiotic activity of SXT under anaerobic conditions, the *rexBA* mutants were still more susceptible than the corresponding WT, with >10 and 2.5-fold greater zone sizes for JE2 and SH1000 strains respectively (Fig. 3B).

**Figure 3.**
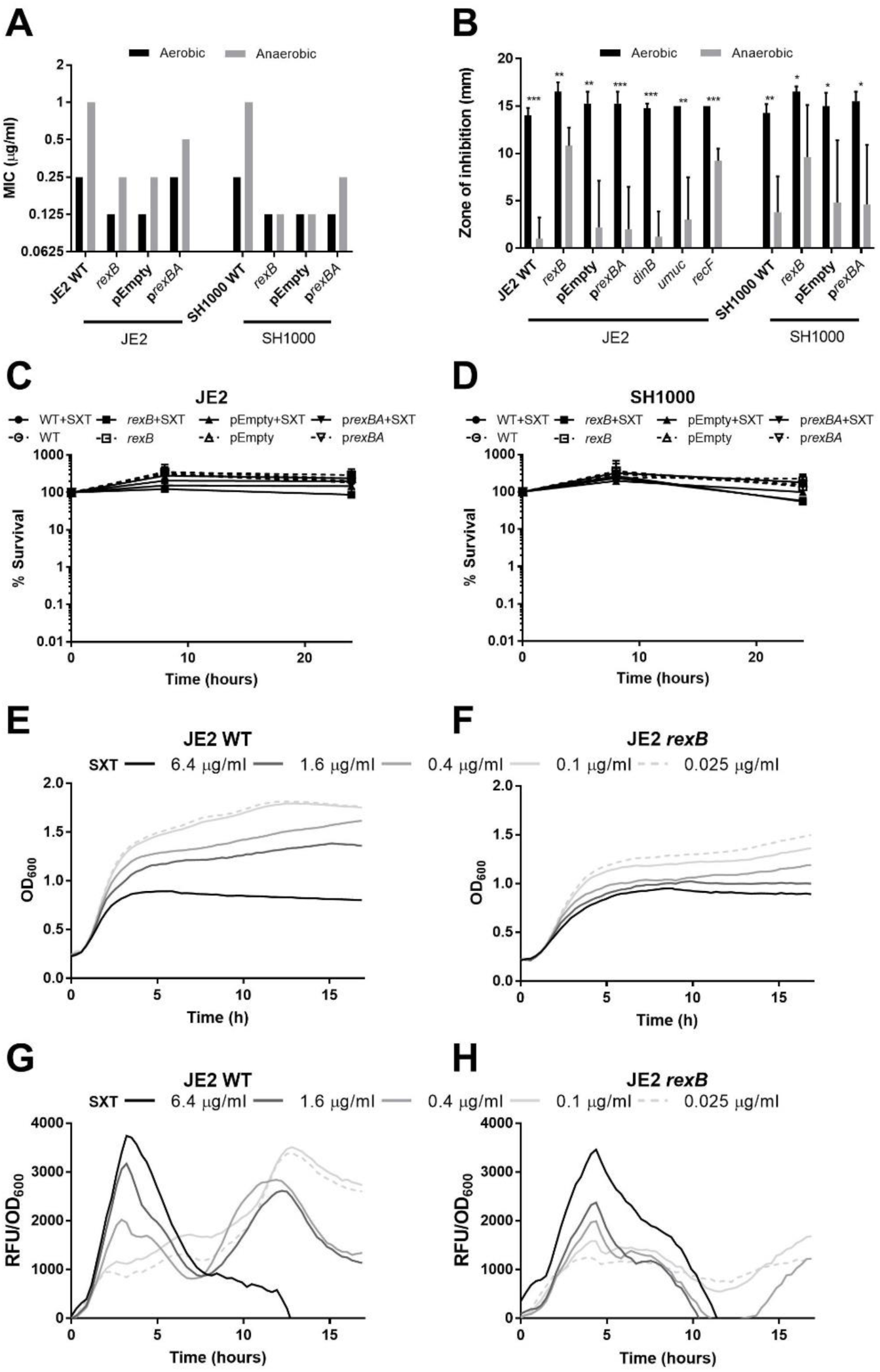
Reactive oxygen species are essential for SXT-mediated killing of *S. aureus*. (A) MIC of SXT for WT *S. aureus* JE2 and SH1000 and their derived mutants defective for DNA repair determined in TSB under aerobic or anaerobic conditions. Graphs represent median values from n=3. (B) Zones of inhibition around paper discs containing SXT (2.5 μg) on agar plates inoculated with *S. aureus* JE2 WT or SH1000 WT and associated DNA repair mutants on TSA after 16 h of incubation under aerobic (black) or anaerobic (grey) conditions. The graph represents mean±s.e.m from n=4. Zones which are significantly different in size between aerobic and anaerobic conditions were determined by a two-tailed Student’s t-test. (C,D) Time-course survival assays of *S. aureus* JE2 (C) and SH1000 (D), *rexB* mutants without or with empty vector (pEmpty) or complemented (p*rexBA*) in TSB with (+SXT) or without 4 μg ml^-1^ SXT, under anaerobic conditions. Graphs represent mean±s.e.m from n=3. Values that are statistically different from the WT were determined by two-way ANOVA corrected for multiple comparison using the Sidak method. P *≤* 0.05(*), P *≤* 0.01(**), P *≤* 0.001(***), P *≤* 0.0001(****). (E-F) Detection of growth inhibition (OD_600_) of WT JE2 (E) and *rexB* mutant (F) in TSB in the presence of SXT. (G-H) Detection of ROS production by bacteria using the DCF fluorophore for *S. aureus* JE2 WT (G) or *rexB* mutant (H) in the presence of SXT. RFU generated by ROS were normalised to OD_600_ data (E,F) to determine ROS production relative to cell density. (E-H) Error bars were omitted for clarity.

The elevated susceptibility of the *rexB* mutant to SXT under both aerobic and anaerobic conditions indicated that thymidine limitation causes double-strand breaks via a process that is promoted by, but not dependent upon, the presence of oxygen.

Next, we assessed the contribution of ROS to the bactericidal activity of SXT. In contrast to assays done in air, none of the strains examined were killed by SXT (4 μg ml^-1^) under anaerobic conditions (Fig. 3C,D). However, whilst the WT strains was able to grow slightly under these conditions, SXT prevented growth of the *rexB* mutants >2-fold by 24 h post inoculation (Fig. 3C,D). The ability of the WT strains to grow in the presence of an SXT concentration above the MIC likely reflects the 1000-fold larger inoculum used in time kill experiments compared to MIC assays (36-38). Complementation of the *rexB* mutant, but not the empty vector, rescued SXT mediated growth inhibition for both strains at 24 h (Fig. 3C,D). Therefore, SXT exposure causes bacteriostasis in the *rexBA* mutant but not the WT under anaerobic conditions.

The reduced activity of SXT under anaerobic conditions suggested that ROS make an important contribution to the bactericidal activity of the antibiotic. Therefore, H_2_DCFDA dye, which is converted to the fluorescent ‘DCF’ in the presence of ROS, was added to growth inhibition assays (Fig. 3E,F) to quantify and study the kinetics of ROS production during incubation under aerobic conditions. SXT induced clear concentration dependent inhibition growth for both the JE2 WT and *rexB* mutant (Fig. 3E,F). When these data were used to normalise to the relative cell density, SXT also induced a dose-dependent fluorescent peak at 4 h post inoculation for both WT and *rexBA* mutant strains in the JE2 background (Figures 3G and 3H), indicative of the generation of ROS in response to the antibiotic. A second peak in ROS production after 13 h is apparent at lower concentrations of SXT as cells reach stationary phase, which is likely a result of the bacterial starvation response, which is associated with increased endogenous oxidative stress (39).

Combined, these data provide strong evidence that SXT-induces ROS that contribute to the growth inhibitory activity of the antibiotic and are essential for the bactericidal activity of the drug. Furthermore, these findings provide evidence that SXT triggers similar levels of oxidative stress in both WT and *rexB* mutant, indicating that the increased susceptibility of the mutant to the antibiotic is due to an inability to repair damage, rather than increased generation of ROS.

### RexAB is required for SXT-triggered induction of the SOS response

The SOS response is a DNA damage-inducible regulon that includes several genes that encode DNA repair components and is regulated by RecA and LexA (17). Consistent with our findings that SXT causes DNA damage in *S. aureus*, previous work has shown that TMP triggers *recA* expression, which is indicative of induction of the SOS response (14). This is important because the SOS response increases the mutation rate, promoting the likelihood of drug resistance emerging, which is a frequent limitation of SXT therapy (17, 40-42).

Since RexAB was important for the repair of DNA damage caused by SXT, we wanted to determine how the complex affected the induction of the SOS response. To investigate this, we used *S. aureus* JE2 WT and *rexB* mutant strains containing a *recA*-*gfp* reporter construct (p*recA*-gfp) to measure induction of the SOS response (Fig. 4).

**Figure 4.**
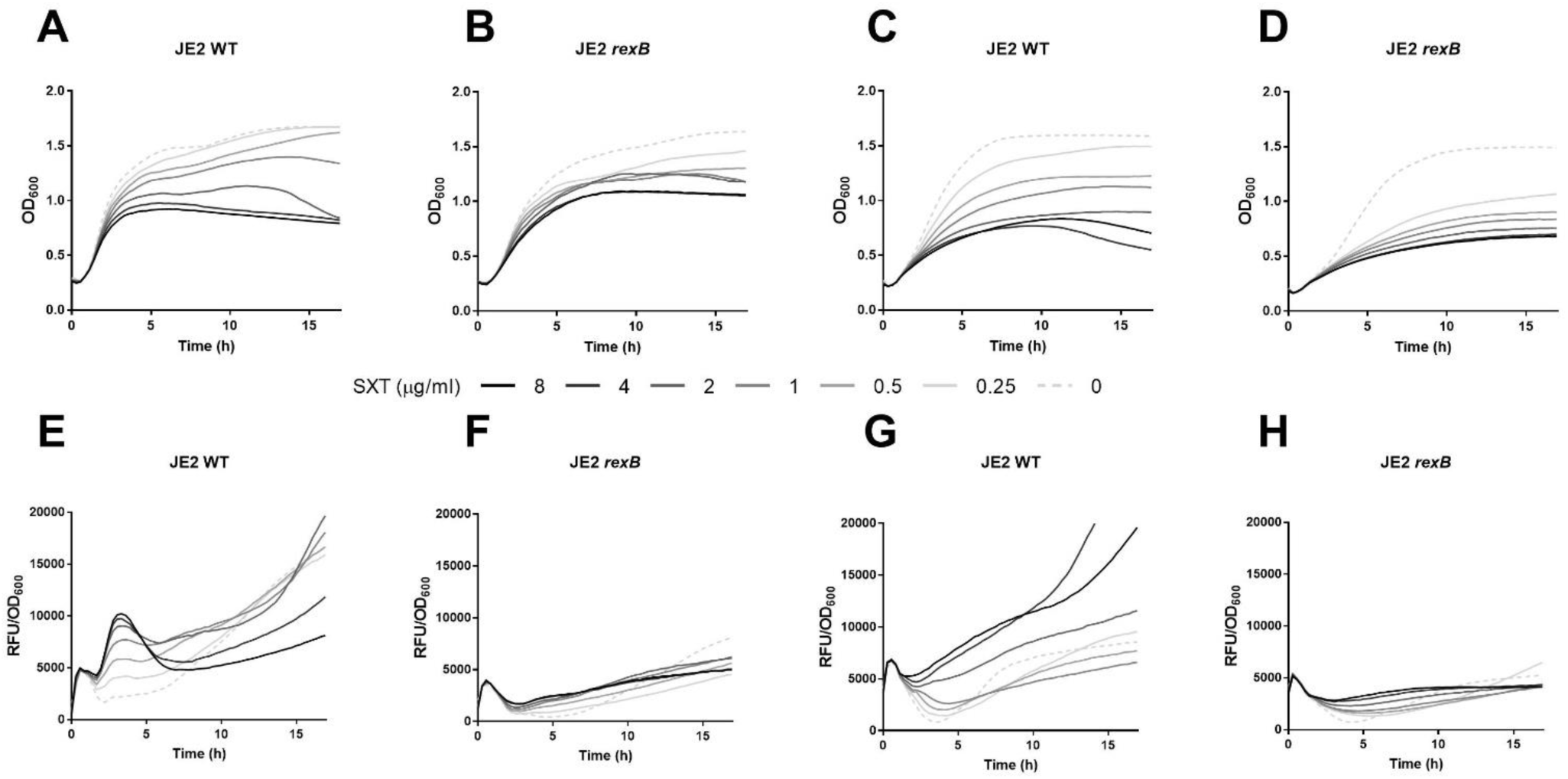
RexAB is essential for the initiation of the SOS response in response to SXT. (A-D) Growth (OD_600_) of WT JE2 (A,C) and *rexB* mutant (B, D) in low thymidine TSB (A,B) and high thymidine MHB (C,D) in the presence of a range of SXT concentrations. (E-H) Expression of *recA-gfp* in response to SXT concentrations in *S. aureus* JE2 WT (E,G), *rexB* mutant (F,H) in TSB (E, F), or MHB (G-H). Fluorescence generated by GFP was measured and normalized to OD_600_ readings to relate *recA* expression relative to cell number. (A-H) Graphs represent mean values from n=3. Error bars were omitted for clarity.

Irrespective of thymidine concentration, SXT induced a dose dependent inhibition of growth for both the JE2 WT and *rexB* mutant (Fig. 4A-D). In low-thymidine TSB, SXT exposure resulted in a dose-dependent increase in GFP signal from the p*recA*-gfp reporter construct in the *S. aureus* WT strain (Fig. 4E), which peaked at about 4 h and then subsided before a the signal slowly increased once more as the bacteria entered stationary phase, presumably due to SOS induction in response to internal DNA-damaging events as a consequence of nutrient limitation and other metabolic stresses (39, 43).

The temporal profile of *recA* expression is similar to that observed for ROS production by WT *S. aureus* in response to SXT (Fig. 3G), providing additional evidence that antibiotic-induced oxidative stress contributes to DNA damage. However, although SXT also triggered ROS production in the *rexB* mutant (Fig. 3H), the antibiotic did not induce *recA* expression in this strain (Fig. 4F), revealing that RexAB is required for activation of the SOS response under these conditions.

In contrast to experiments done in TSB, *recA* was only weakly expressed during exposure to SXT in high-thymidine MHB (Fig. 4D), which is in keeping with the reduced activity of the antibiotic in this medium (Fig. 2E,F). Consistent with the data from studies in TSB, there was a lack of *recA* expression in the JE2 *rexB* mutant in response to SXT in MHB (Fig. 4E).

### RexAB is required for SXT-induced mutagenic DNA repair

De-repression of the SOS regulon leads to expression of the error-prone translesion DNA polymerase UmuC, which catalyses mutagenic DNA repair that contributes to the acquisition of antibiotic resistance (17, 20).

To assess whether SXT promoted SOS-dependent mutagenesis in *S. aureus*, JE2 WT and DNA-damage repair mutants were exposed to sub-inhibitory concentrations of SXT in low-thymidine TSB. The resulting mutation rate was calculated by fluctuation analysis using the emergence of rifampicin resistance, which occurs via point mutations in the *rpoB* gene encoding the RNA polymerase β-subunit (30, 44). Since the *rexBA* mutant was more susceptible to SXT treatment than the wild-type, only a very low concentration of the antibiotic (0.05 µg ml^-1^, 0.2 x MIC of the mutant) could be used in mutation rate analyses with this strain. However, even at this concentration, SXT treatment resulted in an increase in the mutation rate of WT *S. aureus* (Fig. 5A). By contrast to the WT, SXT exposure did not increase the mutation rate of the *rexB* mutant, consistent with the inability of the antibiotic to trigger induction of the SOS response in this strain (Fig. 4B, 5A).

**Figure 5.**
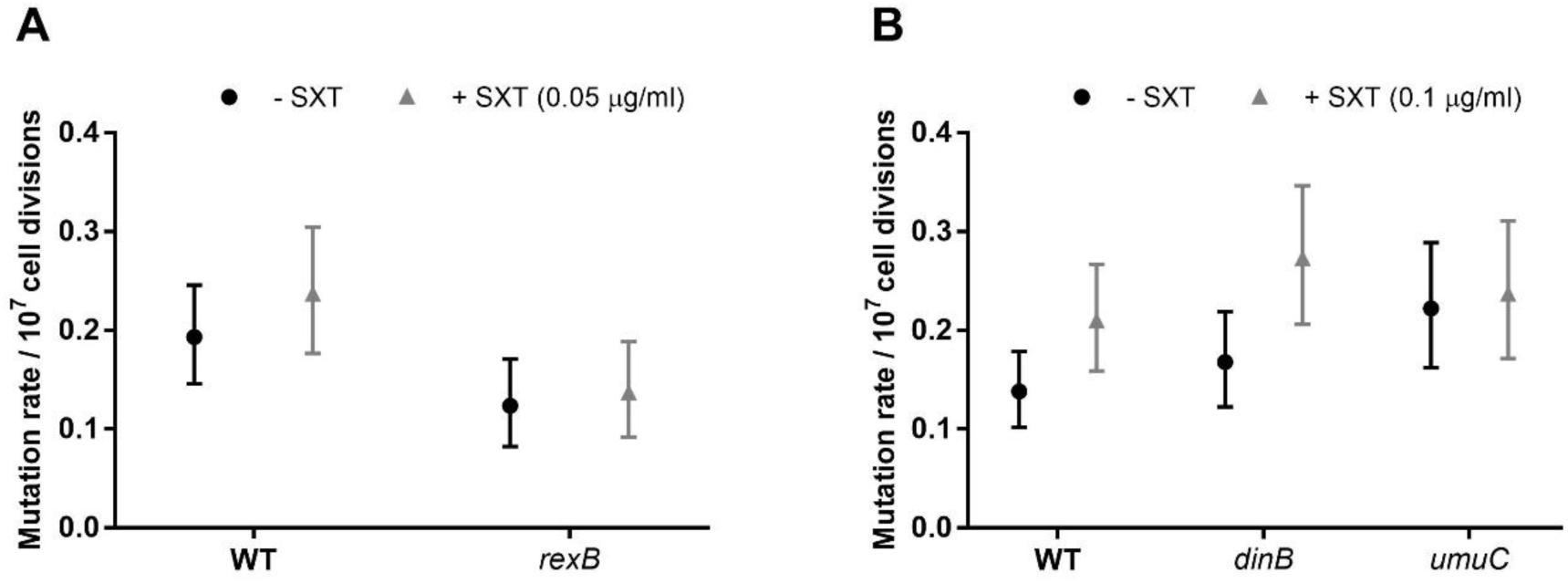
RexAB and UmuC are required for the increased mutation rate mediated by SXT. (A,B) The mutation rate of *S. aureus* JE2 or mutants lacking components of DNA repair in TSB in the absence or presence of 0.25 μg ml^-1^ (A) or 0.5 μg ml^-1^ (B) SXT. Error bars represent 95% confidence intervals.

To determine whether the SXT-mediated increase in the mutation rate was due to induction of the SOS response, we exposed the WT and mutants lacking *umuC* or *dinB* (an error prone polymerase that is not part of the SOS response in *S. aureus*; (17) to SXT at 0.2 x MIC of the mutants (0.1 µg ml^-1^). Consistent with the *recA* reporter data, the higher concentration of SXT used in these experiments promoted the mutation rate of the WT strain to a greater extent than the lower dose (Fig. 4A, 5A,B). SXT caused a similar increase in the mutation rate of the *dinB* mutant but had no effect on the mutation rate of the *umuC* mutant (Fig. 5B).

Taken together, these data demonstrate that SXT exposure results in RexAB-dependent induction of the SOS response in *S. aureus*, which triggers UmuC-dependent mutagenic DNA repair that promotes the rate at which antibiotic resistance arises.

## Discussion

Multi-drug resistance in *S. aureus* is a major global health concern associated with high treatment failure rates (45). Many second or third-line treatments such as vancomycin or daptomycin can lack efficacy, require intravenous administration and are toxic (12). By contrast, SXT is a safe and orally administered antibiotic that is effective in the treatment of skin and soft tissue infections caused by MRSA (4). However, the efficacy of this antibiotic is limited by the pathogens’ ability to by-pass SXT-mediated inhibition of folate biosynthesis, either via the uptake of metabolites from the environment or by the frequent acquisition of resistance-conferring mutations (9). Therefore, it is hoped that an improved understanding of SXT and its mechanism of action will to enable the development of novel approaches aimed at expanding the therapeutic use of this antibiotic and reducing the frequency at which resistance emerges.

Although poorly characterised, the bactericidal effect of SXT has been ascribed to DNA damage, which leads to the induction of the SOS response in bacterial cells (14, 15). The data we present here support DNA damage as a central component of the mechanism of SXT bactericidal activity in *S. aureus* and highlight the key role of RexAB in combatting this genotoxicity. In contrast to the WT, mutants defective for *rexB* were significantly more susceptible to SXT in both growth inhibition and killing assays.

RexAB is proposed to be a member of the AddAB family of DNA repair enzymes required for the processing of DNA double strand breaks on the basis of protein sequence identity (19, 20, 46). Therefore, the increased susceptibility of *rexB* mutants to SXT indicate that DNA DSBs are caused by the antibiotic and are lethal if not repaired. Additional evidence that SXT causes DNA damage in *S. aureus* came from analysis of the *recF* mutant, which was more susceptible to the antibiotic than the WT, albeit to a lesser extent than the *rexB* mutant. The function of RecF has been ascribed to the repair of DNA lesions that arise from DNA single-strand gaps. However, whilst this function is well established in canonical model organisms for bacterial DNA damage repair, including *E. coli* and *Bacillus subtilis*, the evidence that supports RecF performing a similar function in *S. aureus* is less strong (18). Therefore, a greater understanding of how RecF repairs damage in staphylococci will provide insight into the type of DNA damage caused by SXT in *S. aureus*. Whilst we didn’t see increased SXT susceptibility for any of the other mutants examined in MIC assays, this doesn’t imply that other types of DNA damage don’t occur, simply that they don’t contribute to bacterial growth inhibition. This finding does imply however, that DNA repair does not contribute to the lethality of SXT, as has been reported for trimethoprim-mediated killing of *E. coli* (13).

The discovery that *S. aureus* is more susceptible to SXT when incubated in thymidine-poor medium is in keeping with previous reports, which suggest that thymidine depletion makes a crucial contribution to the lethality of SXT (15, 35). Specifically, nucleotide imbalance, caused by thymidine depletion, can result in the collapse of replication forks during DNA synthesis, which can lead to DNA DSBs (8). In support of this, we observed SOS induction under thymidine-limited but not thymidine-replete conditions. This has important implications for the efficacy of SXT-treatment since elevated thymidine concentrations are often observed in human tissues containing necrotic cells or neutrophil extracellular traps (9, 47). To acquire this thymidine, *S. aureus* expresses DNase to break down DNA and NupC, a nucleoside transporter that has been shown to support the growth of *S. aureus* during SXT exposure by enabling the uptake of extracellular thymidine (48, 49). The presence of thymidine and other folate-dependent metabolites including serine, methionine and glycine in host tissues modulate the susceptibility of SXT-treated cells and restrict the types of infection that can be treated with this antibiotic (9). However, whilst the presence of thymidine reduced the susceptibility of the WT to SXT, there was still a very large reduction in CFU counts of the *rexB* mutant exposed to the antibiotic, suggesting that thymidine limitation is not the only element responsible for DNA damage in *S. aureus*.

In agreement with several reports that bactericidal antibiotics, including TMP, exert part of their antibacterial effect via the generation of ROS (13, 50), we determined that the bactericidal effect of SXT against *S. aureus* is dependent upon the availability of oxygen. This was evident by the complete lack of bacterial killing under anaerobic conditions and suggests that ROS-mediated DNA damage makes a greater contribution to the bactericidal activity of SXT than the inhibition of DNA-synthesis caused by thymidine depletion in aerobic conditions.

Antibiotic-induced oxidative stress increases the intracellular level of 8-oxodeoxyguanosine (8-oxo-dG), which is incorporated into DNA, and paired with deoxycytidine or deoxyadenosine (51). Incomplete repair of DNA-incorporated 8-oxo-dG can lead to DSB in *E. coli*, which contributes to the lethality of TMP (13). Additional experiments using mutants defective for the repair of 8-oxo-dG are needed to determine whether a similar mechanism mediates the genotoxicity of SXT in *S. aureus* under aerobic conditions. However, whilst ROS clearly make an important contribution to the bactericidal activity of SXT, the increased susceptibility of the *rexB* mutant to this antibiotic under anaerobic conditions confirms that DNA damage can occur independently of ROS. This may explain why there are phenotypic differences in the effects of SXT exposure and thymineless death observed in *thyA* mutants deprived of thymine (13).

One drawback of SXT is the frequent emergence of resistance during long-term therapy (52). In keeping with the induction of the SOS response, exposure of *S. aureus* to sub-inhibitory concentrations of SXT was found to increase the mutation rate via UmuC, a low-fidelity DNA polymerase, previously shown to mediate mutagenic DNA repair of bacteria exposed to ciprofloxacin or H_2_O_2_ (20). Similar to what has been reported for ciprofloxacin and H_2_O_2_, SXT-induced increases in the mutation rate was dependent upon RexAB (20). Therefore, the processing of DNA DSBs by RexAB is central to the survival of *S. aureus* exposed to SXT as well as SOS-mediated increases in the mutation rate associated with drug resistance.

In summary, our data demonstrate that SXT causes DNA damage in *S. aureus* via thymidine depletion and ROS generation, the repair of which enables bacterial survival and leads to induction of the SOS-mediated increase in the mutation rate and the emergence of drug-resistance. The identification of RexAB as central to both SXT susceptibility and SOS induction suggests that this protein complex is a promising target for novel therapeutic approaches that could improve the efficacy of SXT and reduce the emergence of resistance. Such an approach may also increase the range of MRSA infections that could be treated with SXT.

## Acknowledgements

A.M.E. and R.S.C. acknowledge funding from Shionogi & Co., Ltd. A.M.E. also acknowledges support from the National Institute for Health Research (NIHR) Imperial Biomedical Research Centre (BRC).

K.P.H. is supported by a PhD scholarship funded by a Medical Research Council award to the Centre for Molecular Bacteriology and Infection (MR/J006874/1). All authors acknowledge the provision of strains by the Network on Antimicrobial Resistance in *Staphylococcus aureus* (NARSA) Program: under NIAID/ NIH Contract No. HHSN272200700055C.

The funders had no role in the study design, interpretation of the findings or the writing of the manuscript.

